# In vivo multiphoton microscopy of cardiomyocyte calcium dynamics in the beating mouse heart

**DOI:** 10.1101/251561

**Authors:** Jason S. Jones, David M. Small, Nozomi Nishimura

**Affiliations:** Nancy E. and Peter C. Meinig School of Biomedical Engineering, Cornell University, Ithaca, United States.

## Abstract

We demonstrated intravital multiphoton microscopy in the beating heart in an intact mouse and optically measured action potentials with GCaMP6f, a genetically-encoded calcium indicator. Images were acquired at 30 fps with spontaneous heart beat and continuously running ventilated breathing. The data were reconstructed into three-dimensional volumes showing tissue structure, displacement, and GCaMP activity in cardiomyocytes as a function of both the cardiac and respiratory cycle.

## Main text

In vivo multiphoton microscopy (MPM) was recently demonstrated within the heart of rodent models^1-6^. However, the combined use of intravital MPM and genetically encoded calcium indicators, which has revolutionized brain studies by enabling in vivo measurement of action potentials from individual neurons within a genetically-defined cell population^7-9^, has not been achieved in the heart. As a result, little is known about calcium dynamics at the level of single cardiomyocytes in the in vivo environment. Previous in vivo studies of calcium dynamics utilized wide-field fluorescence imaging that primarily reports surface calcium transients averaged across many cells^10,11^, while cell-resolved calcium imaging has relied on reduced preparations such as ex vivo Langendorf perfusion models, which eliminate both blood flow and cardiomyocyte contraction^12,13^. Here, we demonstrated methods for imaging activity of the genetically encoded calcium indicator, GCaMP6f^7^, in the beating heart within a living mouse with the capability to resolve single cardiomyocytes, measure calcium dynamics as a function of depth into the ventricle wall, and characterize the dependence on both cardiac and respiratory cycles.

In anesthetized, mechanically ventilated mice, we acquired ∼100-μm thick image stacks with 2-μm step size and 50-100 images per plane through a window mounted to a stabilized probe glued to the left ventricle (Fig. 1a and b, Supple. Fig.1) while recording the electrocardiogram (ECG) and ventilator pressure (Fig. 1c). This preparation caused minimal tissue damage (Supple. Fig. 2a) and heart rate was stable throughout the imaging session (Supple. Fig. 2b). High frame rate imaging (30 fps), using resonant scanners, produced clear images in real time throughout the cardiac cycle, with less distortion due to tissue motion than with slower scanning with galvanometers (Supple. Fig. 3). In contrast to previous approaches^2,3^, breathing was not paused during measurement and image acquisition and heart beat were not synchronized. Instead, the effects of breathing and heart beat were decoupled by reconstructing 3D volumes from smaller image segments sorted by both the cardiac and respiratory phases^14^ (Fig. 1d), with the size of bins in phase and position in z adjusted for the needs of the application. For example, reconstructed 3D stacks from a mouse expressing GCaMP6f in cardiomyocytes and with a vascular injection of dye enabled the clear visualization of blood vessels and cardiomyocytes (Fig. 1e), from which the GCaMP signal in individual cardiomyocytes could be extracted throughout the heartbeat (Fig. 1f). Each stack included data from 10% of the cardiac cycle (defined as starting from the R wave in the ECG) and expiration (50-100% of the respiratory cycle defined as starting from inspiration), with frames averaged over 10 μm of depth.

**Figure 1.**
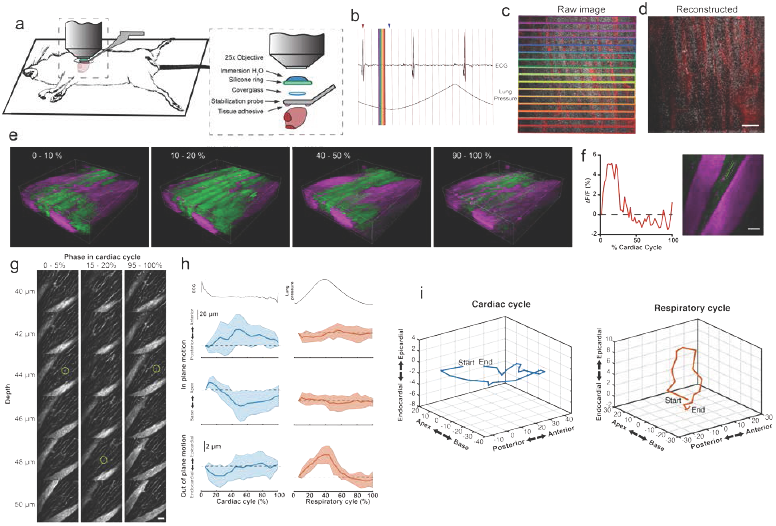
Combination of surgical approach and image reconstruction methods enable visualization and quantification of cardiac cell dynamics. (a) Optical access is gained via a left thoracotomy and a stabilization probe is attached to the left ventricle for microscopy. The probe holds a 3-mm diameter coverglass in contact with the heart. A silicone ring is glued to the top of the probe to hold water for the immersion of the microscope objective. Tissue adhesive is applied on the bottom surface of the probe prior to attachment to the heart. (b) Electrocardiogram (ECG) and ventilator pressure are recorded simultaneously during image acquisition allowing image reconstruction. Red vertical lines indicate the start of each frame; red arrow indicates the peak of R-wave used as the start of the cardiac cycle for the frame displayed below; blue arrow indicates the end of respiratory exhalation that was used as the marker of respiratory cycle. (c) Single raw image frames with colored boxes indicating the image segments, with corresponding timing of the acquisition indicated on the ECG and ventilator pressure traces. (d) A plane reconstructed using 512 x 33 pixel segments, 5% of the cardiac cycle, restricted to 70-100% of the respiratory cycle, and averaged across 4 μm in z. Supplemental Fig. 4 shows intensity as a function of cardiac and respiratory cycle from this data. (e) Renderings of image volumes reconstructed from segments of 512 x 33 pixels, at the indicated cardiac cycle phases, including from 50 to 100% of the respiratory cycle, averaging over 10 μm in depth. GCaMP6 channel shown in green. Intravascular Texas-Red dextran shows vasculature in magenta. (f) GCaMP6f intensity change in a single myocyte throughout the cardiac cycle (∆F/F) indicated in an image plane 50 μm below the heart surface. (g) Representative images of myocardial vasculature labelled with Texas-Red dextran dye used for tracking the 3D motion of a structure (yellow circle) at the indicated phases in the cardiac cycle. Reconstruction used 512 x 65 pixel segments, 5% cardiac cycle bins, included 50-100% of respiratory cycle and used 2-μm bins in z. (h) Average change in the position of features in the myocardium throughout the cardiac cycle from reconstructions from (g) and throughout the respiratory cycle (from same data, reconstructed including data 50–100% of the cardiac cycle, 5% respiratory bins). Solid lines are average and shading indicates standard deviation from 14 measurements from eight stacks in three animals. (i) Three-dimensional average trajectory of the same data showing the average displacement due to cardiac and respiratory motion.

Tissue displacement relative to the stabilized frame was dependent on both cardiac contraction and lung inflation. To quantify this motion throughout the cardiac cycle, point-like structures, such as the edge of a capillary bifurcation, were manually tracked in 3D in reconstructed stacks (Fig. 1g; binned by 5% of the cardiac cycle, averaged over the second half of respiratory cycle). In-plane displacements (base-apex (x) and anterior-posterior (y)) from the initial positions at the start of the cardiac cycle had an elliptical profile and were larger (32.4 ± 31.6 μm, mean ± standard deviation, and 32.3 ± 31.3 μm, respectively; 14 measurements across three animals) than out of plane in the axial (depth in myocardium) direction (3.9 ± 4.7 μm) (Fig. 1h and i), which is consistent with the combination of movements associated with subepicardial helical fiber shortening and ventricle shortening along the longitudinal axis being larger than changes in heart wall thickness^15^. During the respiratory cycle, displacements from the position at lowest ventilation pressure in the base-apex and anterior to posterior direction were small, while structures moved in the ventral direction by 9.5 μm ± 3.8 around the peak lung inflation (Fig. 1j; averaged over second half of cardiac cycle, 5% binning along respiratory cycle). This is likely residual motion due to lung inflation pressing the heart against the stabilized coverglass, and not a respiratory dependent change in thickness of the myocardium.

The GCaMP signal from one imaged volume, normalized to the depth-dependent maximum and averaged through the imaged volume, exhibited a cardiac cycle dependent peak consistent with measurements averaged across regions and individual cells in ex vivo preparations^12^ (Fig. 2a and c) as well as a respiratory cycle dependent decrease (Fig. 2a and d). In this stack, signal from the vascular label decreased shortly after the R-wave and exhibited a respiration-dependent decrease that mirrored that of the GCaMP signal (Fig. 2b, e and f). In wild type mice that did not express the calcium indicator, we imaged autofluorescence using the same emission filters used for GCaMP and observed a small increase in intensity within the first half of cardiac cycle (Supple. Fig. 5a). The change in the fluorescence intensity, however, was about 90 times smaller than in GCaMP-expressing mice. We further measured the GCaMP and vascular label signals across both the respiratory cycle and cardiac cycle as a function of depth into the heart wall (Fig. 2g and h). We calculated rise time and decay time^16^ of the GCaMP signal across the cardiac cycle as a function of imaging depth (11 stacks from three mice, data from 25-50% of the respiratory cycle excluded to avoid confounding effects of the respiratory dependent tissue motion) and found that both rise and decay times remained relatively constant over depths from 15 to 90 μm with mean values of 32 ± 7.4 and 96 ± 36 ms, respectively, and were consistent with other measurements with GCaMP6f^16^ (Fig 2i and j).

**Figure 2.**
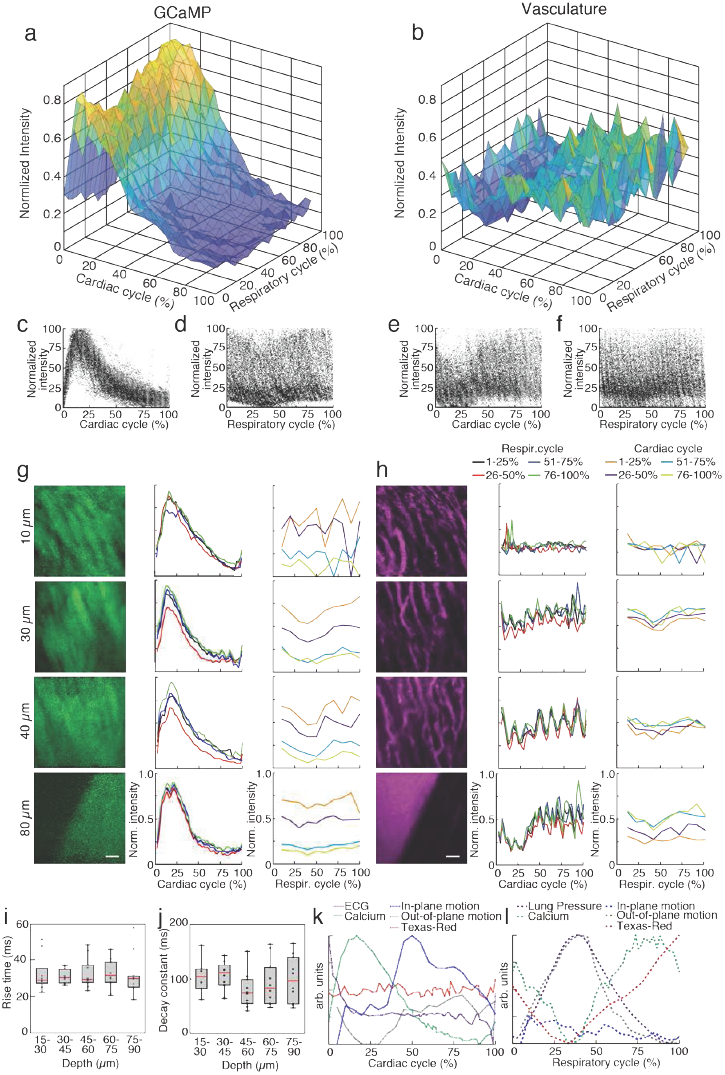
GCaMP imaging in ventricle wall. (a) Average intensity in the calcium indicator GCaMP6f and (b) Texas-Red fluorescence channels as a function of cardiac and respiratory cycle averaged over a stack spanning 90 μm in depth. Displayed values are the average intensities from 512 x 33 pixel segments, normalized to the maximum intensity in the sequential series of raw images acquired at that plane. Data were combined in bins of 25% of both cardiac and respiratory cycle. Plots of the same data showing all individual measurements of GCaMP intensity across (c) cardiac and (d) respiratory cycles and from the Texas-Red channel across (e) cardiac and (f) respiratory cycles. (g) GCaMP channel and (h) Texas-Red channel reconstructed image projections (left), average plots of fluorescence intensities as function of cardiac phase (middle) and breathing phase (right). Images include 50-100% of respiratory cycle, and bin 10% of the cardiac cycle, and include 10 μm in z. The cardiac dependence (middle) used bins of 2% of the cardiac cycle. The respiratory dependence used bins of 10% of the respiratory cycle. Color of lines indicate phase range of trace in opposing cycle. Scale bars are 50 μm. (i) Rise time and (j) decay constants from the calcium responses from 10 regions of interest across three animals (2% bins of the cardiac cycle, excluded 25-50% of the respiratory cycle, and binned 15 μm in depth). Box edges represent 25^th^ and 75^th^ percentiles, box centerline indicated median, and whiskers are most extreme data not considered outlier. ANOVA showed no significant differences in groups (p = 0.7, p = 0.9). (k) Summary of mean trends as a function of cardiac and respiratory phase. Data comes from 10 regions of interest from three animals.

Measurements of motion, action potentials, and Texas-Red fluorescence from the same reconstructed image stacks show correlations with ECG and breathing across the cardiac and respiratory cycles (Fig. 2k and l). Data aggregated across three animals demonstrates GCaMP signal variation with cardiac phase did not correlate with in-plane motion and slightly preceded the out-of-plane motion (Fig. 1h), suggesting that it reflects changes in calcium, rather than artifact from motion (Fig. 2k). The contribution of autofluorescence (Supple. Fig. 5), which is likely dominated by NADH^17,18^, was small compared to the GCaMP signal. The volume of cardiomyocytes stays about constant throughout contraction^19-21^, so that the concentration of GCaMP stays the same and does not contribute to the signal change. Taken together, this suggests that variation in the GCaMP signal is largely attributable to changes in calcium concentration. On average, the vessel fluorescence stayed constant over the cardiac cycle (Fig. 2k), although individual planes showed fluctuations both increasing and decreasing (for example, Fig. 2h) which tended to match timing of the out of plane motion, peaking around 25% of the cardiac cycle. These changes of fluorescence in the vessel channel could come from change in amount of blood plasma in the tissue as vessels are deformed by cardiac pressures. GCaMP and Texas-Red fluorescence, out-of-plane displacement and ventilator pressure all peaked at around 35% of the respiratory cycle, suggesting that GCaMP signal changes with respiration were due to changes in optical properties or residual motion, rather than changes in calcium concentration in cardiomyocytes (Fig. 2l). Both GCaMP and vascular label signals exhibited a decrease in intensity at the peak pressure of inspiration (Fig. 2d and f). Both breathing and heartbeat change the shape and composition of tissue so that future studies will need to take the resulting changes in path length, absorption, scattering, and wavefront distortion into account.

Our methods enable simultaneous measurement and visualization of multiple aspects of cardiac physiology at the cellular scale, including motion, action potentials, and vasculature. The time resolution of the methods allows data collection even during dynamic contraction of the cardiomyocyte. Allowing the animal to respire during measurements enables studies of the coupling between respiratory and cardiac function and facilitates studies of disease models that could be confounded by breath holds. In this method, there is a tradeoff between 1. the degree of binning in cardiac and respiratory phase and in the extent of spatial averaging and 2. the amount of time spent imaging to acquire sufficient data. We demonstrated that 1.5-3s of image acquisition per axial plane is sufficient for characterization across both respiratory and cardiac cycles, so experiments involving stacks before and after a manipulation or time lapse studies are quite feasible. While faster imaging modalities such as ultrasound^22^ and OCT^23^ can measure tissue motion and blood flow more conveniently that MPM, fluorescence techniques have the advantage of the availability of many functional indicators such as GCaMP and labels such as promoter-driven reporter gene expression. Combined with our protocol for in vivo imaging, this makes multiphoton microscopy a potent tool for cardiac studies.

## Methods

Methods, including statements of data availability and any associated references are available in the online version of the paper.

## Acknowledgements

We would like to thank Michael Kotlikoff for advice on cardiac physiology, Chris Schaffer and Chris Xu for sharing optical expertise, Nathan Ellis for machining expertise, Saif Azam and Jui Pandya for contributing to experiments, and George Calvey for assistance with fs laser machining. Funding was provided by the American Heart Association (13SDG17330004 to NN, 17POST33680127 to DMS), Congressionally Directed Medical Research Program (PR151579P1), New York State Department of Health (DOH01-C32240GG-3450000).

## Author Contributions

JSJ and DMS designed and executed experiments, conducted data analysis, and wrote the manuscript. NN designed experiments and wrote the manuscript.

## SUPPLEMENTARY FIGURES

**Supplemental Figure 1.**
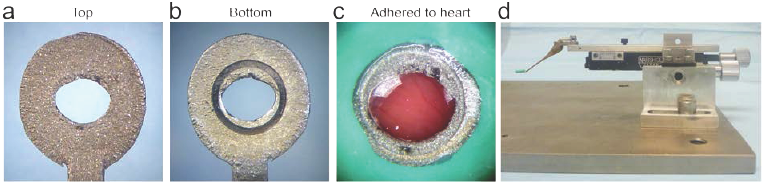
Stabilization probe. (a) top and (b) bottom views of the 3D-printed titanium stabilization probe. A femtosecond laser ablated groove on the underside that prevents tissue adhesive from leaking underneath the coverglass. (c) Probe attached to heart showing attached green silicone ring that serves as reservoir for the microscope objective immersion fluid. (d) Stabilization probe attached to the micomanipulator which allows accurate placement of the probe onto the ventricle wall.

**Supplemental Figure 2.**
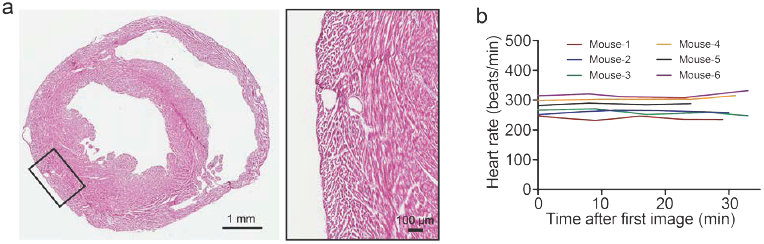
Histology of the imaged heart. (a) Hematoxylin and eosin staining of in vivo, MPM imaged heart demonstrated no observable damage to cell structure (insert). (b) Heart rate during imaging remains constant.

**Supplemental Figure 3.**
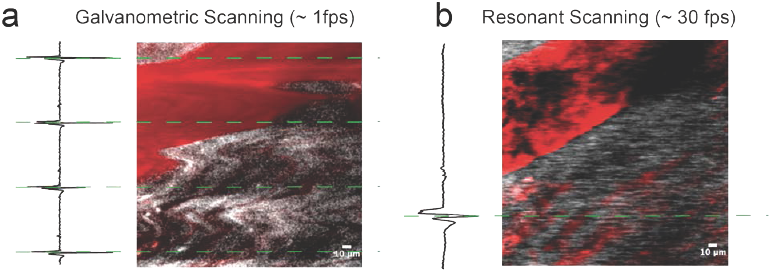
Higher frame rate imaging shows reduced in-frame motion due to heart contraction. Raw image frames showing same cardiac vessel with (a) standard galvonometric scanning and (b) resonant scanning. Green dotted lines indicate the timing of the peak of the R-wave from the electrocardiogram which align with image artifacts. Scale bar is 10 μm.

**Supplemental Figure 4.**
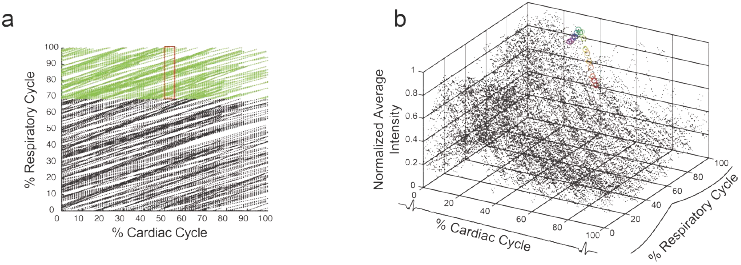
Reconstruction of images with sorting by cardiac and respiratory cycle. (a) Data density from an image stack from Fig. 1b-d with 50 frames acquired per z-plane, 90 μm in z shows coverage of the 2D parameter space. Each point represents a 512 x 33 pixel segment from the raw image data. Figure 1c and d show images from this data set. Green dots indicate the included range of respiratory cycle and the red box indicates portion of the cardiac cycle used to generate the reconstructed plane in Fig. 1d. (b) From the same stack, GCaMP signal as a function of cardiac and respiratory cycles. Each data point is the intensity averaged in a 512 x 33 pixel segment with values normalized to the maximum intensity per axial slice. Circled points correspond to data taken from regions outline in corresponding colors in the raw image frame in Figure 1c.

**Supplemental Figure 5.**
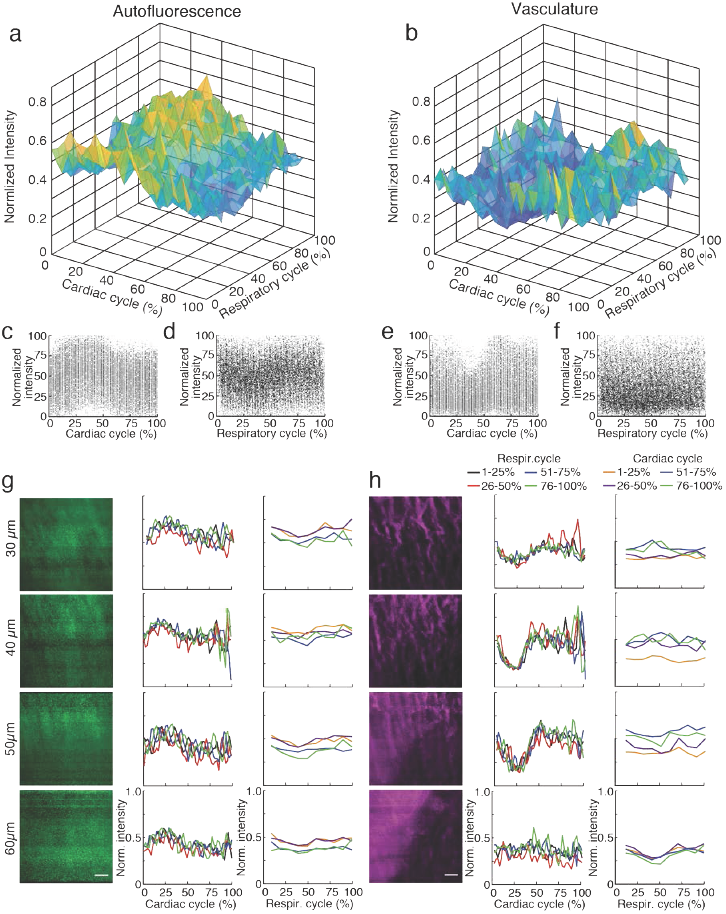
Intensity as a function of respiratory and cardiac cycle in a wild type mouse analyzed in the same manner as Figure 2. Autofluorescence intensity measured with the same wavelength range as for GCaMP6f (517/65 nm filter) and (b) Texas-Red fluorescence as a function of cardiac and respiratory cycle over a stack spanning 90 μm in depth. Displayed values are the average intensities from 512 x 33 pixel segments, normalized to the maximum intensity in the sequential series of raw images acquired at that plane. Data were combined in bins of 25% of both cardiac and respiratory cycle. Plots of the same data showing all individual measurements of autofluorescence intensity across (c) cardiac and (d) respiratory cycles and from the Texas-Red channel across (e) cardiac and (f) respiratory cycles. (g) GCaMP channel and (h) Texas-Red channel reconstructed image projections (left), average plots of fluorescence intensities as function of cardiac phase (middle) and breathing phase (right). Images include 50-100% of respiratory cycle, and 10% of the cardiac cycle, and include 10 μm in z. The cardiac dependence (middle) used bins of 2% of the cardiac cycle. The respiratory dependence used bins of 10% of the respiratory cycle. Color of lines indicate phase range of trace in opposing cycle. Scale bars are 50 μm.

## ONLINE METHODS

### Mice

Mice expressing the cre-dependent GCaMP6f fast variant calcium indicator gene (B6;129S-Gt(ROSA)26Sor^tm95.1(CAG-GCaMP6f)Hze^/J – Jackson Labs; #024105)^7^ were bred in house with B6.Cg-Tg(CAG-cre/Esr1*)5Amc/J (CAG-cre/Esr1) mice (Jackson Labs; #004682)^24^. Expression of GCaMP5f was induced by intraperitoneal injection of tamoxifen in corn oil (Sigma #T5648) for 5 consecutive days (75 mg/kg body weight), resulting in widespread tissue expression including in the cardiomyocytes. C57BL/6 wild-type mice were used for control experiments. Male and female mice (25 – 40 g), aged 4 – 10 months old were used for experiments. All animal care and experimental procedures were approved by the Institutional Animal Care and Use Committee of Cornell University and comply with the Guide for the Care and Use of Laboratory Animals by the National Institutes of Health.

### Design of stabilization window and surgery for imaging

The design of the stabilization window and imaging setup is shown in Figure 1a and b and Supple. Fig.1. The probe is composed of 3D-printed titanium (iMaterialize, Belgium) and the head contains a 2-mm central aperture that accepts a 3-mm diameter coverslip (Deckglaser Cover Glasses, 64-0726 #0). The tissue interface side of the probe was sanded and a channel was etched around the central aperture using fs laser ablation with 1-kHz, 15-μJ, 800-nm, 50-fs laser pulses, translated at 0.1 mm/s and focused with a 0.28 NA microscope objective. The groove around the circumference of the underside of the aperture prevents tissue adhesive spilling underneath the coverslip and impairing image quality. A silicone ring, fashioned from silicone molding putty (Castaldo Qick-Sil), is adhered with Loctite-406 to the top side of the window to hold water for immersion of the microscope objective at the appropriate working distance. The tail of the probe is fixed to a micromanipulator that can translate the height of the window facilitate placement onto the ventricle wall, and provides an anchorage point for stabilization during imaging.

### Surgical preparation

Mice were anesthetized with ketamine (10 mg/mL) and xylazine (1 mg/mL) in saline via intraperitoneal injection (0.1 mL/10g body weight) and then intubated via the trachea using a 22-gauge, 1-inch catheter to allow mechanical ventilation (95 breath/min, 12 cm H_2_O end-inspiratory pressure; CWE Inc. SAR-830/P ventilator) with medical grade oxygen and 1.5% isoflurane for maintaining anesthesia. The ventilation rate was selected for optimum animal stability and also to avoid harmonics of the heart rate so that time of R wave would cycle through different phases of the respiratory cycle in several seconds. The mouse was positioned on its right side, on top of a motorized stage with a heating pad to maintain body temperature at 37.5 °C. Hair over the left thorax was depilated, skin and muscle layers over the chest wall excised, and the intercostal space between ribs 7 and 8 perforated and retracted to create space for placement of the window. Following removal of the pericardial sac, stabilization probe was attached to the left ventricle free-wall using tissue adhesive (Vetbond). Tissue adhesive was applied to the underside of the probe and the window was gently lowered onto the left ventricle wall. A three-axis micromanipulator, mounted on the motorized stage holding the animal, provided stabilization and positioning of the probe (Fig. 1). Electrocardiogram (ECG) electrodes were attached to 21 gauge needles inserted subcutaneously through the front and contralateral hind limb, connected through an isolated differential amplifier (World Precision Instruments; #ISO-80), and recorded inside a Faraday enclosure mounted to the microscope. ECG and ventilation pressure signals (from ventilator) were continuously monitored on an oscilloscope and recorded simultaneously with image acquisition. 5% glucose in saline (0.1 mL/10 g body weight) and 0.15 mg/mL atropine sulfate (5 μg/100 g body weight) was injected subcutaneously every 30 min throughout surgery and in vivo microscopy. This procedure allowed stable in vivo imaging for 1 h. A retro-orbital injection of Texas-Red conjugated, 70 kDa dextran (3% in saline; Thermo #D1830) was performed to label blood plasma providing contrast in the vasculature.

### In vivo cardiac multiphoton microscopy

Imaging was conducted using a custom multiphoton microscope equipped with four detection channels and an 8-kHz resonant scanner (Cambridge Technology) imaged onto a galvanometric pair (Cambridge Technology) to allow for both resonant and line scanning. Resonant scanning data acquisition was performed using a National Instruments digitizer (NI-5734), FPGA (PXIe-7975), and multifunction I/O module (PXIe-6366) for device control, mounted in a PXI chassis (PXIe-1073) controlled by ScanImage 2016b. A Ti:Sapphire laser (Chameleon, Coherent) with the wavelength centered at 950 nm, was used to simultaneously excite GCaMP6f and Texas-Red fluorescence. Emission was separated using a primary dichroic (Semrock FF705-Di01-49x70), a secondary dichroic (Semrock FF593-Di03-90x108), and bandpass filters selective for GCaMP6f (517/65) and Texas-Red (629/56). Water was placed within the silicone O-ring of the stabilization probe to immerse the microscope objective (Olympus XLPlan N 25x 1.05 NA). ECG and respiratory voltage signals were collected with the two unused detection channels allowing simultaneous recording during imaging. Z-stacks of images to a depth of 100–150 μm from the surface (2 μm per z-step) were collected at 4 – 5 different locations per mouse. A series of 50–100 frames (1.7 to 3.3 s) per plane in z were collected at a scan speed of 30 frame/sec.

### Assigning cardiac and respiratory phase to image

The R-wave and expiration peaks were found by using *findpeak*, a Matlab signal processing toolbox function, to generate a lookup table for cardiorespiratory-dependent image reconstruction and quantification. We found that with a heart rate of about 5 Hz and breathing at 2 Hz, ∼1.5 seconds or about 50 frames was sufficient to generate images in most of the cardiac/respiratory cycle phase space (Supple. Fig.4a and b), although longer imaging periods could be useful for more averaging and to guarantee sampling across the full space. Raw images were divided into segments of *n_block_* lines in the fast-scan axis (x) direction. Because the scan rate in x-direction (0.058 ms per line) is fast compared to the dynamics of the cardiac cell, we assigned image segments a single value of time relative to heartbeat and breathing. For all image segment, we generate a look-up table with position in y, position in z, time relative to the preceding R wave, time relative to the preceding trough of pressure, average intensity values, raw time from preceding R-wave and pressure trough, a unique identifier of each heartbeat, which was were used for transient and image reconstruction. Phase in cardiac cycle was defined as

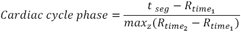

and phase in respiratory cycle was defined as

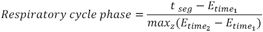

where *t*_*seg*_ is the time of a given segment, *R*_*time*_1__ is the time of the preceding R wave, *R*_*time*_2__ is the time of the following R wave, *E*_*time*_2__ is the time of the preceding trough of pressure, *E*_*time*_2__ is the time of the following trough of pressure, and *max*_*Z*_ is the maximum occurring at a given z position. For exclusion of vascular regions in the quantification of fluorescence, the cardiomyocytes were identified by manually setting intensity thresholds to ignore pixels outside of the cells.

### Reconstruction of image stack

Image planes and volumes are reconstructed by sorting segments by cardiac and respiratory phase. Each line within a selected range of cardiac and respiratory phases was then registered by the position in y and z into a reconstructed stack. In cases where more than one image segment registered to the same position in the reconstruction, the redundant data was averaged. In many applications, segments with a range of z-positions can be averaged together if the loss in spatial resolution is acceptable. The choice of segment size, and binning in the cardiac and respiratory cycles affect the temporal and spatial resolution of the method. The binning parameters and the amount of data acquired should be varied to fit the needs of the experiment.

### Calculation of rise time and tau

Intensity values from stacks were averaged over 15 μm in depth. To eliminate non-calcium dependent signals coming from non-cellular autofluorescence, data from the top 15-30 μm of the raw image stack was not included so that the analysis used only frames with visible calcium transients. Due to the drop in fluorescent signal associated with inspiration, data from 25-50% of the respiratory cycle was excluded. Intensity was binned across 2% of the cardiac cycle and normalized to the maximum intensity in the series of images acquired at a particular depth. A single term exponential curve was fit from 90% of the peak intensity of the transient to the minimum for calculation of tau^16^. The average time from the R wave to the maximum value of the transient was used for the rise time.

### Image display and rendering

ImageJ (imagej.nih.gov/ij) was used to display and process reconstructed images. A median filter (1 pixel) was applied and contrast adjusted for display using only linear scaling. When displaying changes in intensity of GCaMP6f, the displayed range of values was the same at each time point. Renderings of 3D stacks were produced using Imaris x64 version 9.0.2 (Bitplane). The blend mode in Imaris was used to display the volume following the application of a smoothing filter (3x3x1 pixel size).

### Single-point motion tracking

To quantify motion in the myocardium due to cardiac contraction, images were reconstructed by averaging over 5% of the cardiac cycle time, only using images captured during 50–100% of the respiratory cycle. To quantify motion in the myocardium due to respiratory movement, images were reconstructed averaging over 5% of the respiratory cycle, only using images captured during 50–100% of the cardiac cycle. Both cardiac and respiratory cycle reconstructions used 65 x 512 segments of each frame. ImageJ was used to display reconstructed stacks combined at each 2 μm depth (represented in Fig 1f), with each frame of the stack representing sequential 0–5% averaged images throughout the cardiac or respiratory cycle. This allows the point of a small vessel bifurcation to be tracked throughout each 0–5% averaged increment of the cycle and each position (*x, y, z)* to be recorded. These measurements were repeated across three mice consisting of a total of 8 image stacks and of 14 tracked single points.

### Software and code

Matlab was used for reconstruction and cardiac/respiratory phase-dependent analysis. Scripts are available in Supplement Materials. Matlab was used for box plots and statistics analysis.

### Histology

At the end of imaging the probe was removed and the animal deeply anesthetized, followed by transcardial perfusion with cold (4°C) phosphate buffered saline (PBS, pH 7.4, Sigma-Aldrich) followed by 4% (w/v) paraformaldehyde (PFA, ThermoFisher Scientific) in PBS. The heart was excised and cut in the cross-section plane at the level of probe attachment which was indicated by remnants of tissue adhesive. Hearts were placed in 4% PFA for 1 day, and then in 30% sucrose (w/v) in PBS for 1 day. The heart was frozen in Optimal Cutting Temperature (OCT) compound (Tissue-Tek) and cryo-sectioned at 7-μm thickness onto glass slides. Hematoxylin and eosin staining was performed using standard procedures.

## REFERENCES

1. Li, W. et al. Intravital 2-photon imaging of leukocyte trafficking in beating heart. The Journal of clinical investigation 122, 2499–2508 (2012).

2. Aguirre, A.D.,Vinegoni, C.,Sebas, M. & Weissleder, R. Intravital imaging of cardiac function at the single-cell level. Proc Natl Acad Sci U S A 111, 11257–11262 (2014).

3. Vinegoni, C.,Lee, S.,Aguirre, A.D. & Weissleder, R. New techniques for motion-artifact-free in vivo cardiac microscopy. Front Physiol 6, 147 (2015).

4. Lee, S. et al. Real-time in vivo imaging of the beating mouse heart at microscopic resolution. Nat Commun 3, 1054 (2012).

5. Jung, K. et al. Endoscopic time-lapse imaging of immune cells in infarcted mouse hearts. Circ Res 112, 891–899 (2013).

6. Vinegoni, C.,Aguirre, A.D.,Lee, S. & Weissleder, R. Imaging the beating heart in the mouse using intravital microscopy techniques. Nat Protoc 10, 1802–1819 (2015).

7. Chen, T.W. et al. Ultrasensitive fluorescent proteins for imaging neuronal activity. Nature 499, 295–300 (2013).

8. Prevedel, R. et al. Fast volumetric calcium imaging across multiple cortical layers using sculpted light. Nat Methods 13, 1021–1028 (2016).

9. Pologruto, T.A.,Yasuda, R. & Svoboda, K. Monitoring neural activity and [Ca2+] with genetically encoded Ca2+ indicators. J Neurosci 24, 9572–9579 (2004).

10. Tallini, Y.N. et al. Imaging cellular signals in the heart in vivo: Cardiac expression of the high-signal Ca2+ indicator GCaMP2. Proc Natl Acad Sci U S A 103, 4753–4758 (2006).

11. Jaimes, R., 3rd et al. A technical review of optical mapping of intracellular calcium within myocardial tissue. Am J Physiol Heart Circ Physiol 310, H1388–1401 (2016).

12. Rubart, M.,Wang, E.,Dunn, K.W. & Field, L.J. Two-photon molecular excitation imaging of Ca2+ transients in Langendorff-perfused mouse hearts. Am J Physiol Cell Physiol 284, C1654–1668 (2003).

13. Wasserstrom, J.A. et al. Variability in timing of spontaneous calcium release in the intact rat heart is determined by the time course of sarcoplasmic reticulum calcium load. Circ Res 107, 1117–1126 (2010).

14. Santisakultarm, T.P. et al. In vivo two-photon excited fluorescence microscopy reveals cardiac‐ and respiration-dependent pulsatile blood flow in cortical blood vessels in mice. Am J Physiol Heart Circ Physiol 302, H1367–1377 (2012).

15. Sengupta, P.P.,Tajik, A.J.,Chandrasekaran, K. & Khandheria, B.K. Twist mechanics of the left ventricle: principles and application. JACC Cardiovasc Imaging 1, 366–376 (2008).

16. Hammer, K.P.,Hohendanner, F.,Blatter, L.A.,Pieske, B.M. & Heinzel, F.R. Variations in local calcium signaling in adjacent cardiac myocytes of the intact mouse heart detected with two-dimensional confocal microscopy. Front Physiol 5, 517 (2014).

17. Zipfel, W.R. et al. Live tissue intrinsic emission microscopy using multiphoton-excited native fluorescence and second harmonic generation. Proc Natl Acad Sci U S A 100, 7075–7080 (2003).

18. Blinova, K.,Combs, C.,Kellman, P. & Balaban, R.S. Fluctuation analysis of mitochondrial NADH fluorescence signals in confocal and two-photon microscopy images of living cardiac myocytes. J Microsc 213, 70–75 (2004).

19. Rodriguez, E.K. et al. A method to reconstruct myocardial sarcomere lengths and orientations at transmural sites in beating canine hearts. Am J Physiol 263, H293–306 (1992).

20. Sonnenblick, E.H.,Ross, J., Jr.,Covell, J.W.,Spotnitz, H.M. & Spiro, D. The ultrastructure of the heart in systole and diastole. Chantes in sarcomere length. Circ Res 21, 423–431 (1967).

21. Leu, M.,Ehler, E. & Perriard, J.C. Characterisation of postnatal growth of the murine heart. Anat Embryol (Berl) 204, 217–224 (2001).

22. Cikes, M.,Sutherland, G.R.,Anderson, L.J. & Bijnens, B.H. The role of echocardiographic deformation imaging in hypertrophic myopathies. Nat Rev Cardiol 7, 384–396 (2010).

23. Srinivasan, V.J. et al. Quantitative cerebral blood flow with optical coherence tomography. Opt Express 18, 2477–2494 (2010).

24. Hayashi, S. & McMahon, A.P. Efficient recombination in diverse tissues by a tamoxifen-inducible form of Cre: a tool for temporally regulated gene activation/inactivation in the mouse. Dev Biol 244, 305–318 (2002).

